# Unveiling Novel Biocontrol Strategies: Serratia marcescens chiA Gene Against Myzus persicae

**DOI:** 10.1101/2024.06.29.601316

**Authors:** Ahmet Can, Ömür Baysal

## Abstract

**Background:** Microorganisms produce a diverse array of enzymes with potential applications in biological control and pest management. Among these enzymes, chitinase stands out due to its safety for non-target organisms and the environment. Chitinase enzymes play a crucial role in breaking down chitin, which serves as a critical component in insect exoskeletons and fungal cell walls. As a result, they emerge as valuable tools for managing both agricultural pests and pathogens.

**Results:** The chiA gene region of *Serratia marcescens* GBS19 was successfully amplified via PCR and cloned into expression vectors. The resulting chiA protein was expressed and purified through His-tag affinity chromatography. The purified chiA enzyme exhibited optimal activity at 40 °C and pH 5. The insecticidal properties of the purified chiA enzyme were tested against the agricultural pest *Myzus persicae*, revealing an LD_50_ value of 15,804 ppm. Comparative analysis with ref_seq chiA enzymes demonstrated that our purified enzyme shows 98.93% similarity, indicating a high degree of conservation and likely functional similarity. Bioinformatics modelling highlighted a strong binding affinity (-4.10 kcal/mol) between the enzyme and chitin, which was also confirmed with modelled chitin layer and enzyme interaction.

**Conclusion:** The study underscores the potential of *S. marcescens* GBS19 chitinase as an effective and environmentally safe biocontrol agent. The chiA enzyme exhibits promising insecticidal properties, specifically against *M. persicae*, and its strong binding affinity to chitin supports its effectiveness. Given its safety for non-target organisms and the environment, *S. marcescens* GBS19 chitinase holds significant promise as a tool for integrated pest management, contributing to sustainable agricultural practices using directed recombinant DNA technology.

## 1 INTRODUCTION

Chitin is the most common polymer in nature after cellulose and is produced by organisms at a rate of approximately 10^1^^2^-10^1^^4^ tons per year. It is found in a variety of organisms, including the outer skeletons of marine animals, insect shells, and fungi.^1,2^ Additionally, it can be derived from green algae and yeasts.^3^ The chitin has a cellulose-like structure and consists of β-(1,4)-bound D-glucosamine (GlcNAc) units.^4^ Different crystalline forms of chitin exist; α-chitin is the most common and is found in arthropods, while β-chitin is found in cuttlefish pencil and γ-chitin is also found in *Ptinus* beetle cocoon fibres and the stomach of *Loligo*.^5^ Chitin is insoluble in most organic solvents and water but can be made soluble by conversion to chitosan through the deacetylation process.^6^

Chitinases (EC 3.2.1.14) are enzymes that hydrolytically break down chitin, a polysaccharide consisting of N-acetylglucosamine units, encoded for various purposes in various organisms. These enzymes are found in a variety of organisms, including fungi, plants, insects, crustaceans, and bacteria.^7,8^ Chitinases are classified within families 18 or 19 of glycosyl hydrolases. Bacterial chitinases are typically classified within family 18, whereas some *Streptomyces* chitinases are grouped within family 19. Bacterial chitinases are divided into three groups: A, B, and C. The majority of characterized bacterial chitinases belong to group A.^8,9^ Microbial chitinases are produced by bacteria and viruses and degrade chitin structures in insects, thereby increasing the risk of microbial infection of hosts.^10^ Chitinases can directly degrade chitin into low-molecular-weight chitooligomers. In particular, ChiA enzymes, which are particularly active in high pH environments, have the potential for applications in agriculture, industry, and medicine.^11,12^

*Serratia marcescens* is a gram-negative bacterium that produces a prominent level of chitinase enzyme and is one of the most effective bacteria for chitin degradation. It produces at least three chitinases (ChiA, ChiB, and ChiC) and other enzymes, including chitobiase and CBP21 (a probable chitin-binding protein). The chitinolytic mechanism of *S. marcescens* has been extensively characterized, although it remains unclear whether these proteins represent the full chitinolytic mechanism. Furthermore, it was identified as the most active chitinase producer among 100 organisms tested.^13,14^ Chemical pesticides are a common method of controlling plant-invasive insects. However, chemical pesticides have several disadvantages, including high toxicity, the potential to cause diseases in humans and other organisms, and the damage they can cause to the environment. The depletion of the exoskeleton (cuticle) and peritrophic matrix (PM) in insects can be employed to elucidate the impact of chitinases on insects. Microbial chitinases can be employed as environmentally friendly insecticides, thereby increasing crop yields.^15,16^

*Myzus persicae*, commonly known as the peach aphid, represents a significant economic threat to agricultural and horticultural crops. This aphid causes direct damage by feeding on plant food, as well as indirect damage by carrying numerous virus vectors. A few factors contribute to the significance of this species as a pest, including its distribution, host diversity, the mechanisms by which it causes damage to plants, its life cycle, its dispersal capacity, and its ability to develop resistance to pesticides. *M. persicae* is a highly cosmopolitan species with a worldwide distribution and a host range of more than 400 species in 40 different plant families, including many economically important crop plants.^17,18^

This study was conducted by cloning and characterizing the gene encoding the chiA enzyme (GenBank: WRW24584.1) from our original strain GBS19 (NCBI ID: 11761001) belonging to *S. marcescens.* The bacterial strain was previously isolated from greenhouse soils in the Ortaca province of Muğla.

The objective of the study was to use the purified enzyme as a biological and environmentally friendly solution against *M. persicae*, to offer it for biological control. To the best of our knowledge, there is no study examined the insecticidal activity of the chiA enzyme on *M. persicae*.

### 2 MATERIALS AND METHODS

### 2.1 Bacterial growth and culture conditions

The *S. marcescens* GBS19 strain was cultured on Nutrient Agar (NA; Merck) plates at 37°C and stored at 4°C. To produce chitinase, Nutrient Broth (NB; Merck) containing 0.2% (w/v) colloidal chitin was employed. The cultured sample, which had been grown for 24 hours at 37°C, was agitated at a speed of 180 revolutions per minute (rpm).The chitin powder obtained from blue crab (*Callinectes sapidus*) shells, which was sourced from the fish market of the Dalyan town of Muğla, was used.^19^ A suspension of 2 g of powdered chitin in 40 mL of concentrated HCl was prepared and dissolved using a magnetic stirrer. Following that, 100 mL of distilled water was added to the solution, and the mixture was centrifuged at 5000 g at 4°C for 20 minutes. The pellet, which dissolved in distilled water and was adjusted to a pH above 3.5, served as the colloidal chitin and was stored at 4°C.

### 2.1 **Gene Cloning**

Genomic DNA was isolated from the *S. marcescens* GBS19 strain using the GeneJET Genomic DNA Purification Kit (Thermo Scientific) following the manufacturer’s instructions. This DNA was used as a template for PCR. Primers were designed for the amplification of the chiA gene region which was previously submitted to NCBI under the accession Nr. GenBank: WRW24584.1 within the full genome sequence of the GBS19 isolate. These primers were named forward primer (SM-GBS19 Fw: 5’- CCCGGATCCATGCGCAAATTTAATAAACCGCTG-3’) and reverse primer (SM-GBS19 Rv: 5’-CCCAAGCTTTGAACGCCGGCGCTGTT-3’). To facilitate cloning, the BamHI cleavage site was incorporated into the forward primer, while the HindIII restriction site (underlined) was included in the reverse primer. A total volume of 50 μL was used for amplification, comprising 25 μL of High Fidelity 2X Master Mix (Ampliqon), which contained 1 μL (10 μM) of forward primer, 1 μL (10 μM) of reverse primer, and 1 μL (50 ng) of genomic DNA. The reaction was initiated with an initial denaturation temperature of 94°C for 3 minutes and continued for 35 cycles of 1-minute denaturation at 94°C, 1-minute annealing at 65°C, 2 minutes of extension at 72°C, and the final extension was performed for 10 minutes at 72°C. The resulting PCR product was then run on a 0.8% (w/v) agarose gel and purified using the GeneJET PCR Purification Kit (Thermo Scientific). This was followed by ligation into the pBluescript II KS (+) (Stratagene) vector using T4 DNA Ligase (Thermo Scientific). Finally, transformation into *Escherichia coli* One Shot™ TOP10 Electro competent (Thermo Scientific) strain was performed. A selection of plasmids containing the GBS19 chiA gene region was conducted in Petri dishes containing ampicillin (100 μg/mL) and X-gal (20 mg/mL). Recombinant plasmids were isolated using the GeneJET Plasmid Miniprep Kit (Thermo Scientific). To ascertain the accuracy of the chiA gene region, digested fragments were sequenced as procurement of services at Sugenomics Co. (Ankara, Turkey). The chiA gene region was extracted from pBluescript II KS (+), purified using the GeneJET Gel Extraction Kit (Thermo Scientific), and subjected to restriction with BamHI and HindIII enzymes. The truncated chiA fragment was ligated into the truncated pET-22b (+) (Novagen) expression vector with the same enzymes. The resulting plasmids were transformed into the *E. coli* BL21 (DE3) (Gold Biotechnology) strain, and the recombinant plasmids were selected in the presence of AMP (100 μg/mL).

### 2.2 Protein expression and purification

The resulting transformants were planted in 250 mL LB medium containing AMP (100 μg/mL) and incubated at 225 rpm at 37°C until the OD_600_ value reached 0.5-0.6. Once the desired density had been reached, the expression of the chiA protein was induced at 30°C by the addition of 0.5 mM isopropyl-β-D-thiogalactopyranoside (IPTG), and the samples were incubated for four hours. Subsequently, the samples were centrifuged at 5500 x g for 15 minutes at 4°C. Approximately 1 g of the resulting pellet was resuspended on ice using 3 mL of 1X LEW Buffer. Lysozyme was added to a final concentration of 1 mg/mL and incubated in a mixer on ice for 30 minutes. Subsequently, the samples were subjected to ultrasonication in an ultrasonic water bath for ten minutes, with a fifteen-second interval for cooling on ice. The resulting crude lysate was then centrifuged at 10,000 x g for 30 min at 4°C. The supernatant was then transferred into a clean Eppendorf tube, ensuring that the pellet was not disturbed.

The columns were equilibrated with 1X LEW Buffer and allowed to drain by gravity. The supernatant was then added to the equilibrated column. The column was then washed twice with 320 μL of 1X LEW Buffer. Subsequently, 1X Elution Buffer was introduced into the column, and the polyhistidine-tagged chiA protein was transferred to a new Eppendorf tube. The Protino Ni-TED 150 Packed Columns (Machery-Nagel) were employed for these processes. The purified protein was quantified using bovine serum albumin (BSA; Thermo Scientific) as the standard according to the Bradford method.^21^ The size of the chiA protein was confirmed and visualized by 12% sodium dodecyl sulfate (SDS)-polyacrylamide gel electrophoresis (PAGE).

### 2.3 Enzyme activity and optimization of pH and temperature

The chitinase activity was determined by the spectrophotometric method using the chitin azure (Sigma) substrate. A total of 50 μL of the chitin azure solution (5 mg/mL in McIlvain buffer, pH 5.2) and an equal volume of the phosphate buffer solution (pH 7) were added to an Eppendorf tube. The quantity of enzyme was then added to the mixture at a concentration of 2 μg, and the mixture was incubated in a water bath at a temperature of 40°C for 30 minutes. Subsequently, the reaction mixture was centrifuged at 13,000 × g for 5 minutes to terminate the enzyme reaction. The absorbance values of supernatant was measured at 560 nm according to the blank (chitin azure + buffer). The average value was determined according to middle values of triplicate samples. An increase of 0.01 in the absorbance value was considered to be equivalent to one unit (U) of the chitinase enzyme.^22^ All measurements were conducted using 96-well plates, with the Multiskan™ GO Microplate (Thermo Scientific) employed for data acquisition. To assess the influence of pH on the activity of the chitinase enzyme, citric acid buffer (pH 3.0-5.0), phosphate buffer (pH 6.0-8.0), and glycine buffer (pH 9.0-10.0) solutions were employed. To ascertain the optimal pH value for the enzyme, the spectrophotometric method was employed to quantify enzyme activity. The effect of temperature on enzyme activity, measurements were recorded at temperatures ranging from 30°C to 100°C, with an increase of 10°C between each measurement. A volume of 50 μL of the pH 5 buffer in which the enzyme exhibited optimal activity was combined with an equal volume of the substrate. Subsequently, 2 μg of the chiA enzyme was added and incubated for 30 minutes at the indicated temperatures. The activities were quantified at a wavelength of 560 nm.

### 2.4 Insecticidal activity tests on *Myzus persicae*

The *M. persicae* pest was obtained for in-vivo efficiency tests from the Faculty of Agriculture, Plant Protection Department in the Malatya Turgut Özal University, with the kindly delivery of Assoc. Prof. M. Keçeci. To maintain the laboratory production of aphids, eggplant leaves were infected to test peach aphids on untreated plant leaves, eggplant seedlings were grown for three weeks before contamination to remove any insecticides they may contain (Supporting Information Fig. S1). The weevils were collected from the infested leaves using a brush (Supporting Information Fig. S2). The activity of the purified chiA enzyme on *M. persicae* was determined according to the method 001 recommended by the Insecticide Resistance Action Committee (IRAC). The leaves were cut with a disk to a diameter of 3 cm. The leaf discs were immersed in different enzyme concentrations (130 ppm, 100 ppm, 50 ppm, 40 ppm, 20 ppm, and 10 ppm) for 10 seconds, after which they were left on damp blotting paper to dry (with the bottom side facing upwards). The control was conducted using pure water (Supporting Information Fig. S3). Subsequently, 2% agar was prepared in a petri dish with a diameter of 6 cm, and leaf discs immersed in enzyme concentrations were placed in the petri dish with the bottom side of the leaf facing up. Subsequently, five *M. persicae* adults were placed in each petri dish and allowed to grow for 24 hours. After 24 hours, the adults were collected from the leaves using a brush. The petri dish was then covered with stretch film, which was pierced with a needle to allow air to flow. The Petri dishes were incubated at a temperature of 22 ± 1°C and under a 16/8-hour light/dark cycle. The leaves in the petri dish were incubated for 72 hours until they were infested with approximately 15 lice. After the designated period, the lice in the Petri dish were identified as either alive or dead using a hand lens. Lice that did not react when touched with a brush and that could not move in a coordinated manner were considered to be dead. This study was carried out in three repetitions. A dose-response analysis was conducted using the IBM SPSS program to determine the LD_50_ estimate for the tested chiA enzyme. The mortality rates were evaluated by applying the Abbott formula to correct the mortality percentage.^23,24^

### 2.5 Bioinformatic analyzes

A modeling study conducted using Swiss-Model revealed that the structural similarities between the *S. marcescens* chiA enzyme registered in the PDB database (PDB DOI: 1EHN) and the chiA enzyme of *S. marcescens* GBS19 (GenBank: PP083416) were 98.7%. To create the model, a reference model was constructed for the chiA enzyme of GBS19, utilizing the *S. marcescens* chiA enzyme from the PDB database as the structural foundation. The reference model precisely reflects the amino acid sequence of the chitinase produced by GBS19. The created model was employed as a fundamental instrument to comprehend the structure of chitinase generated by the chiA enzyme of GBS19 and to investigate its interaction mechanisms. The model was converted to PDB data and employed in modeling studies to monitor protein-ligand interactions. To determine the interactions of the ChiA enzyme with chitin, its substrate, at the molecular level, a model was created by homology matching using the relevant sequences on S. marcescens GBS19. In the model created the data of the chitin molecule in mol2 format was used as the ligand to gain a deeper understanding of the interactions between the ChiA enzyme and the chitin molecule. Subsequently, the coupling room coordinates were determined. A blind docking process was conducted using X, Y, and Z coordinates to gain insight into the interactions between the chitin molecule and the chiA enzyme. A blind docking process was employed to identify potential binding sites between the chiA enzyme and the chitin molecule. The deployment process was conducted at two distinct central locations. The first set of coordinates was X = -1.8, 68.6, -5.7; Y = 56.1, 169.0, 42.7; Z = 27.2, 118.8, 18.5. The second set of coordinates was X = -2.5, 63.3, -7.8; Y = 69.5, 169.3, 44.2; Z = 33.5, 116.3, 18.2. Finally, the E-DOCK server was employed to assess the precision of this blind docking procedure and visualized using PYMOL. This process enabled the verification of whether the generated model produced biologically meaningful and accurate results. The amino acid similarity of the chiA protein of the *S. marcescens* GBS19 isolate was compared with the reference sequence chiA proteins in the NCBI database. The BLAST results indicated that the amino acid sequences of the 10 most similar proteins were recorded. The recorded sequences were subsequently aligned and visualized using the Jalview program. Concurrently, the amino acid sequences and the chiA protein of the isolate in question were employed to construct a phylogenetic tree utilizing the Neighbour-Joining method via the Geneious program.

## 3 RESULTS

### 3.1 Gene Cloning

The genomic DNA of *S. marcescens* GBS19 was analysed for its 260 nm and 280 nm absorbance values and concentration, which were found to be 1.82 and 60 ng/uL, respectively. This sample was selected as the PCR template to amplify the chiA gene. The amplified chiA gene region was analysed by agarose gel electrophoresis and was determined to be 1708 bp in size, including the primers (Fig. 1.). The PCR products were subsequently subjected to enzymatic digestion with BamHI and HindIII restriction enzymes, followed by ligation. Subsequently, the cloning vector containing the gene was transferred to competent *E. coli* TOP10 bacteria by electroporation. To be success in transformation, a plasmid devoid of the gene region was also transformed. The colonies were selected based on their colouration, with those that were white and positive for the chiA gene being isolated for further analysis (Supporting Information Fig. S4). Afterwards, the recombinant plasmids contained the chiA gene, was run into electrophoresis after using restriction endonucleases. Furthermore, the plasmid was linearised using BamHI and loaded onto the gel to facilitate comparison of the bands. The results of the gel electrophoresis demonstrated the successful ligation of the chiA gene into the cloning vector and transformation of the vector into competent bacteria (Fig. 2.).

**Figure 1.**
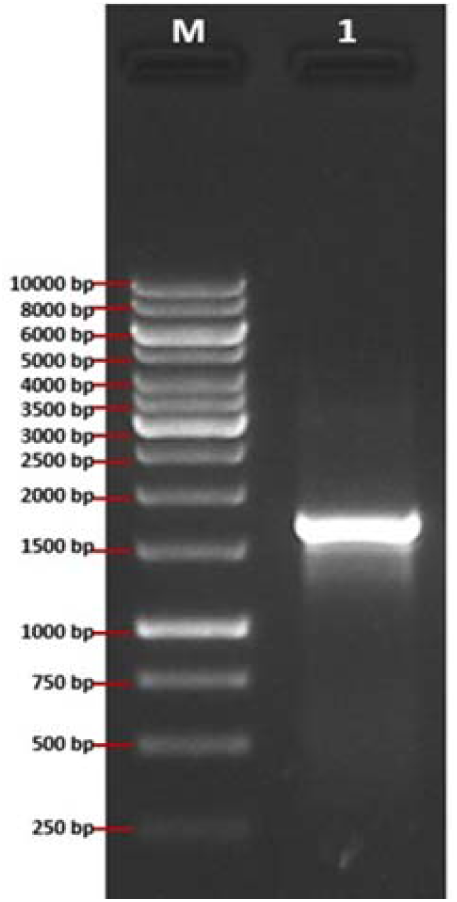
Marker GeneRuler 1 kb DNA ladder (SM0311), 1) PCR result chiA gene region

**Figure 2.**
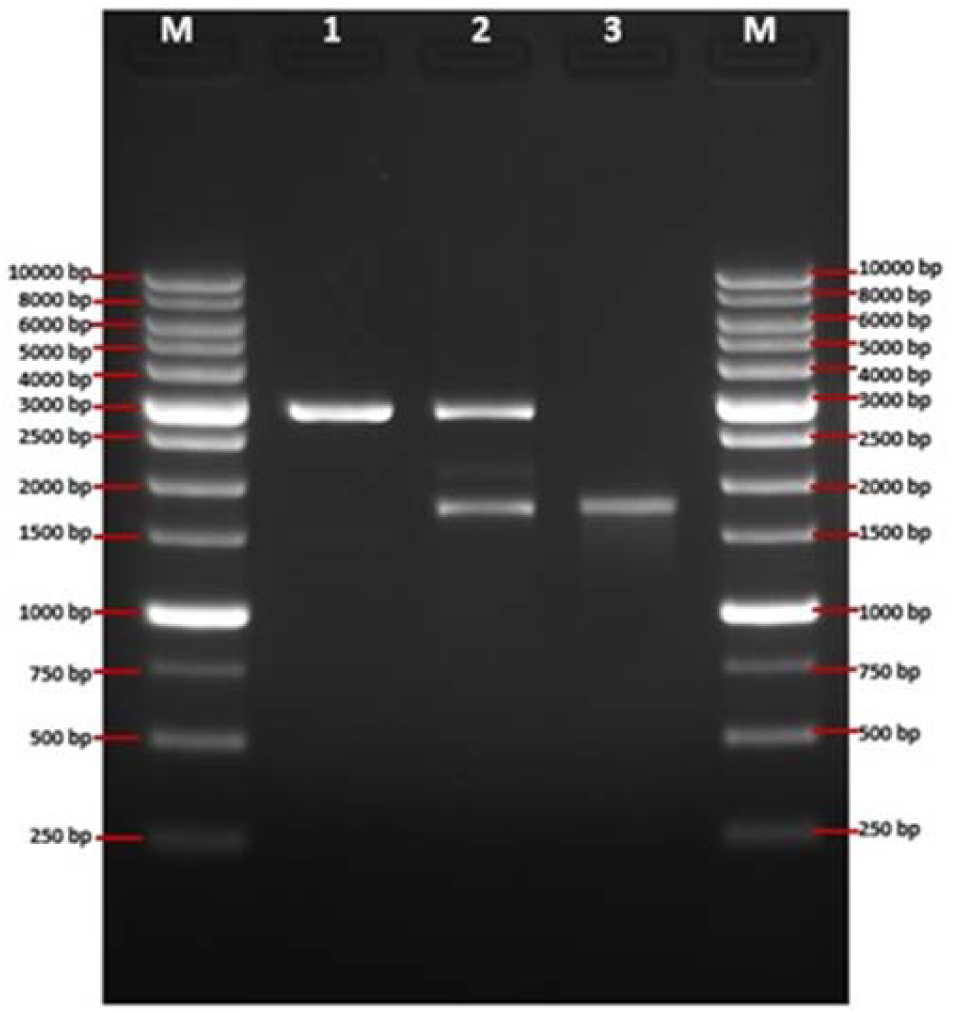
Marker Goldbio 1 kb DNA ladder (D010-500), 1) Single cut with BamHI pBluescript II KS (+), 2) Double cut of recombinant plasmid with HindIII-BamHI, 3) Cut and purified chiA gene region

Sequence analysis of the chitinase gene region contained in the cloning vector confirmed the accuracy of the results (Supporting Information A). The sequence results were visualized using the SnapGene application (Supporting Information Fig. S5). The gene sequences was compared using Jalview program. The cloned chiA gene region exhibited 100% compatibility with the gene sequence obtained through sequencing (Supporting Information Fig. S6).

In order to isolate the chiA gene region from plasmids, the cloning vector was treated with BamHI and HindIII restriction enzymes. The ChiA gene region was subsequently visualised in agarose gel electrophoresis and isolated. Similarly, the pET-22b (+) vector, which was to be used for expression, was cut and cleaned with the same enzymes.

Ligation was performed at a vector/gene ratio of 1:3, and the resulting product was transformed into electrocompetent *E. coli* BL21(DE3). Following the transformation procedure, the bacteria were seeded on LB solid media containing AMP and incubated at 37°C overnight. The plasmid DNA was extracted from the colonies that had been generated following the incubation period. The plasmids contained the chiA gene, double restriction was performed with HindIII and BamHI. Following double cleavage, the chiA gene region was correctly located in the expression vector (Fig. 3.). Concurrently, the recombinant plasmid was sequenced and visualized using the SnapGene software (Supporting Information Fig. S7).

**Figure 3.**
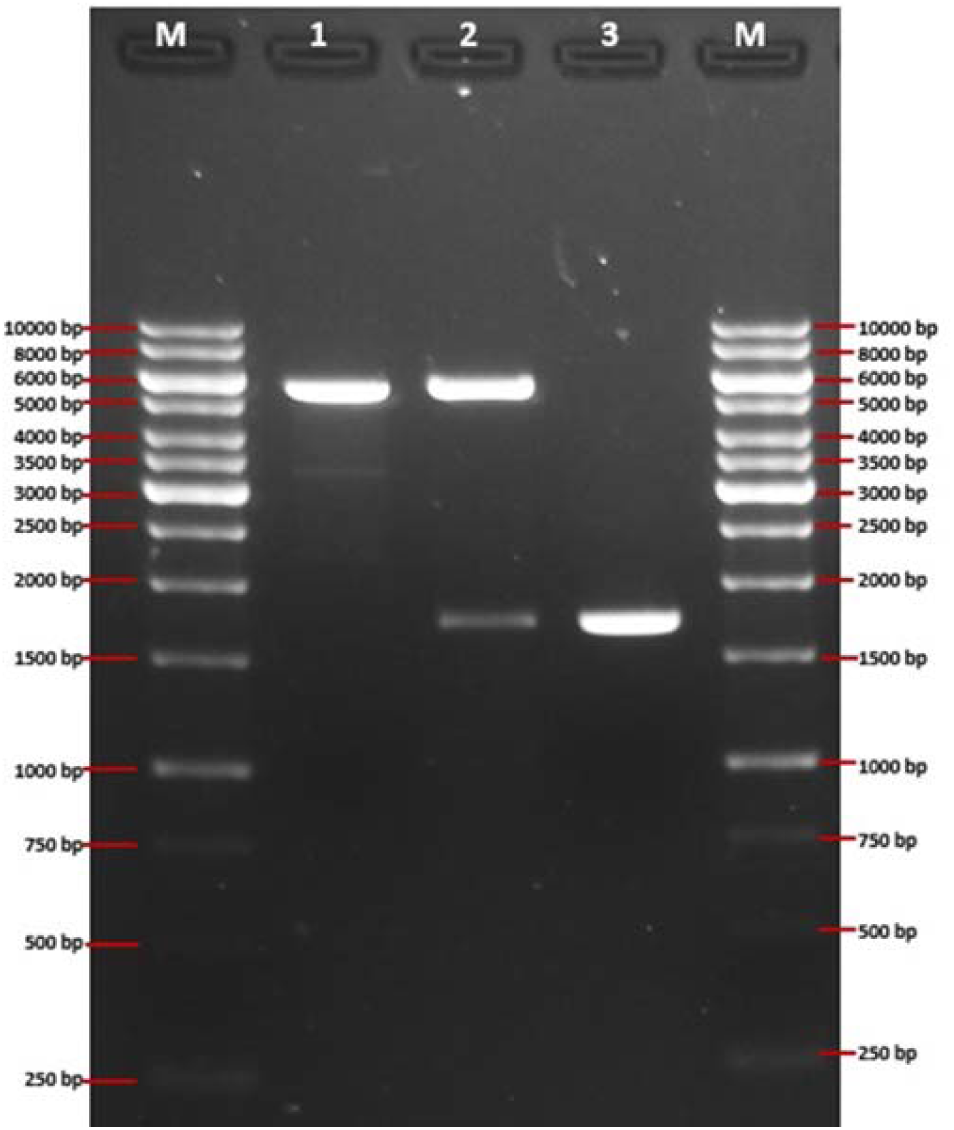
Marker GeneRuler 1 kb DNA ladder (SM0311), 1) Single cut of pET-22b (+) with HindIII, 2) Double cut of recombinant pET-22b (+) with HindIII-BamHI, 3) Cut and purified chiA gene region

### 3.2 Protein production and purification

The *E. coli* BL21(DE3) host cells, which contained the recombinant pET-22b (+) plasmid, were induced with IPTG. The *E. coli* BL21(DE3) strain contains the λDE3 lysogen, which expresses T7 RNA polymerase under the control of the lacUV5 promoter. Upon induction by IPTG, T7 RNA polymerase is expressed.^25^ The pET-22b (+) expression vector also contains the T7 promoter, thus enabling high-level expression. Following the expression of the chiA protein, purification was achieved through the use of Ni-NTA affinity chromatography, utilising the his-tag present within the pET-22b(+) vector. The purified protein was subjected to SDS-PAGE analysis. The analysis yielded the expected result, with the chiA protein exhibiting a 62.5 kDa band on the gel (Supporting Information Fig. S8).

### 3.3 Enzyme activity and optimum pH-temperature

Prior to establishing the optimal pH and temperature values, the optimization was done for the activity of the chitinase enzyme. Trial was also conducted at pH 7 level buffer using Chitin-azure in order to ascertain the activity of the enzyme. The activity experiment carried out with triplicates on the spectrophotometer at a wavelength of 560 nm was calculated according to the absorbance values (Supporting Information Table. T1). An increase of 0.01 in the absorbance value obtained was considered to be equivalent to one unit (U) of chitinase enzyme. The activity of the enzyme was quantified by averaging the absorbance values. In experiments conducted using buffers between pH 3 and 10, the impact of chiA enzyme activity on pH was quantified. Each experiment was repeated three times and measurements were made using a spectrophotometer. The chiA enzyme exhibited the highest activity in a pH 5 buffer (Supporting Information Fig. S9). To determine the effect of chitinase activity on temperature, enzyme activity was measured in experiments performed in a water bath at temperatures between 30°C and 100°C. Each trial was conducted for 30 minutes, with the experiments being repeated on three separate occasions. The mean values of the absorbance were calculated. The optimal temperature for the S. marcescens GBS19 chiA enzyme was 40°C (Supporting Information Fig. S10).

### 3.4 Insecticidal activity

In the activity experiments conducted using the leaf immersion method, the number of live and dead lice living on the leaf was counted at the end of 72 hours of incubation. The results were then compared by creating a graph (Fig. 4.). The insecticidal activity results were used to calculate the enzyme concentrations (LD_50_) required to kill 50% of *M. persicae*. The pure enzyme LD_50_ value was determined to be 15.804 ppm, while the crude enzyme LD_50_ value was 81.090 ppm. The lethality of the unpurified chiA enzyme was approximately five times less than the pure chiA enzyme.

**Figure 4.**
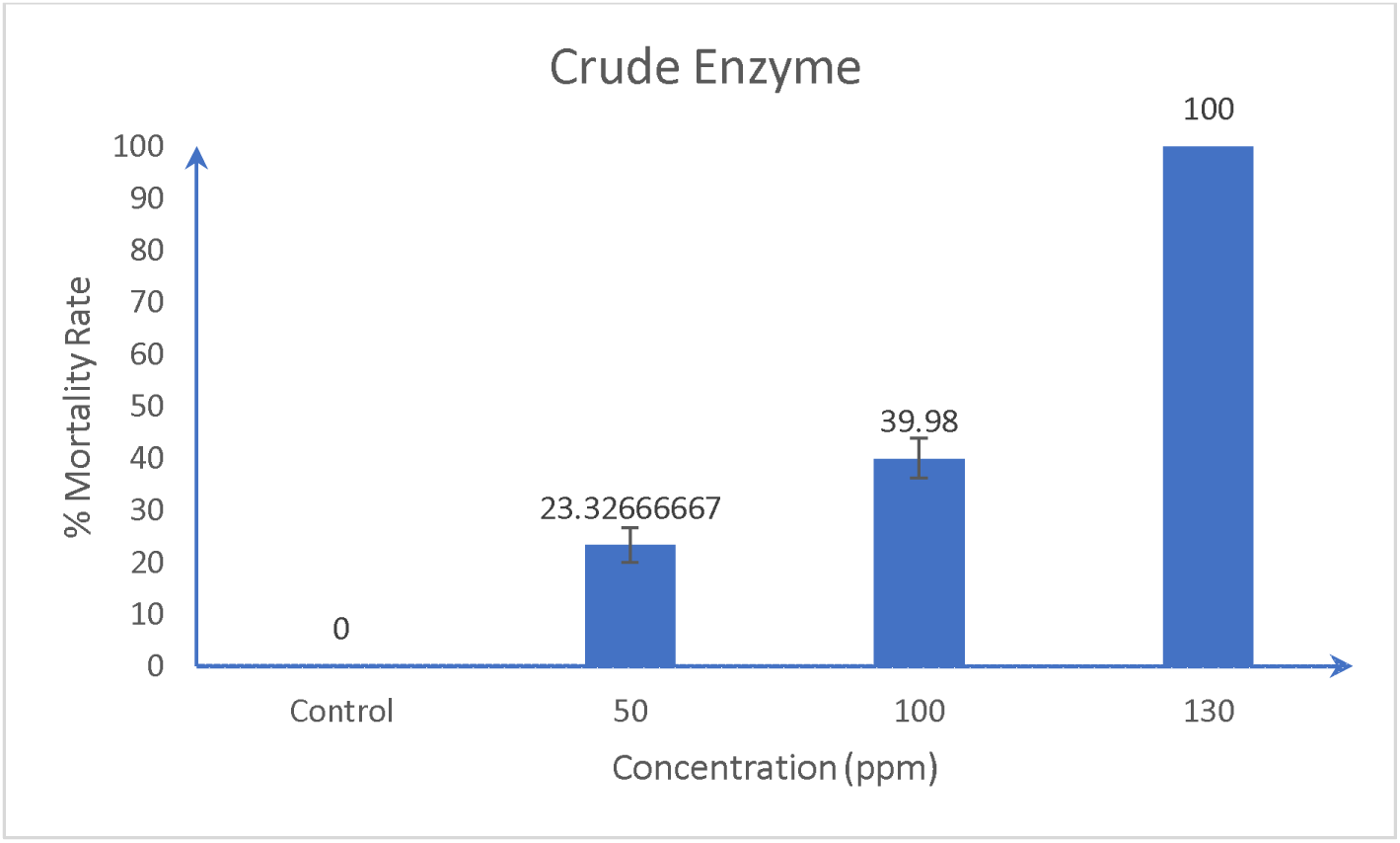
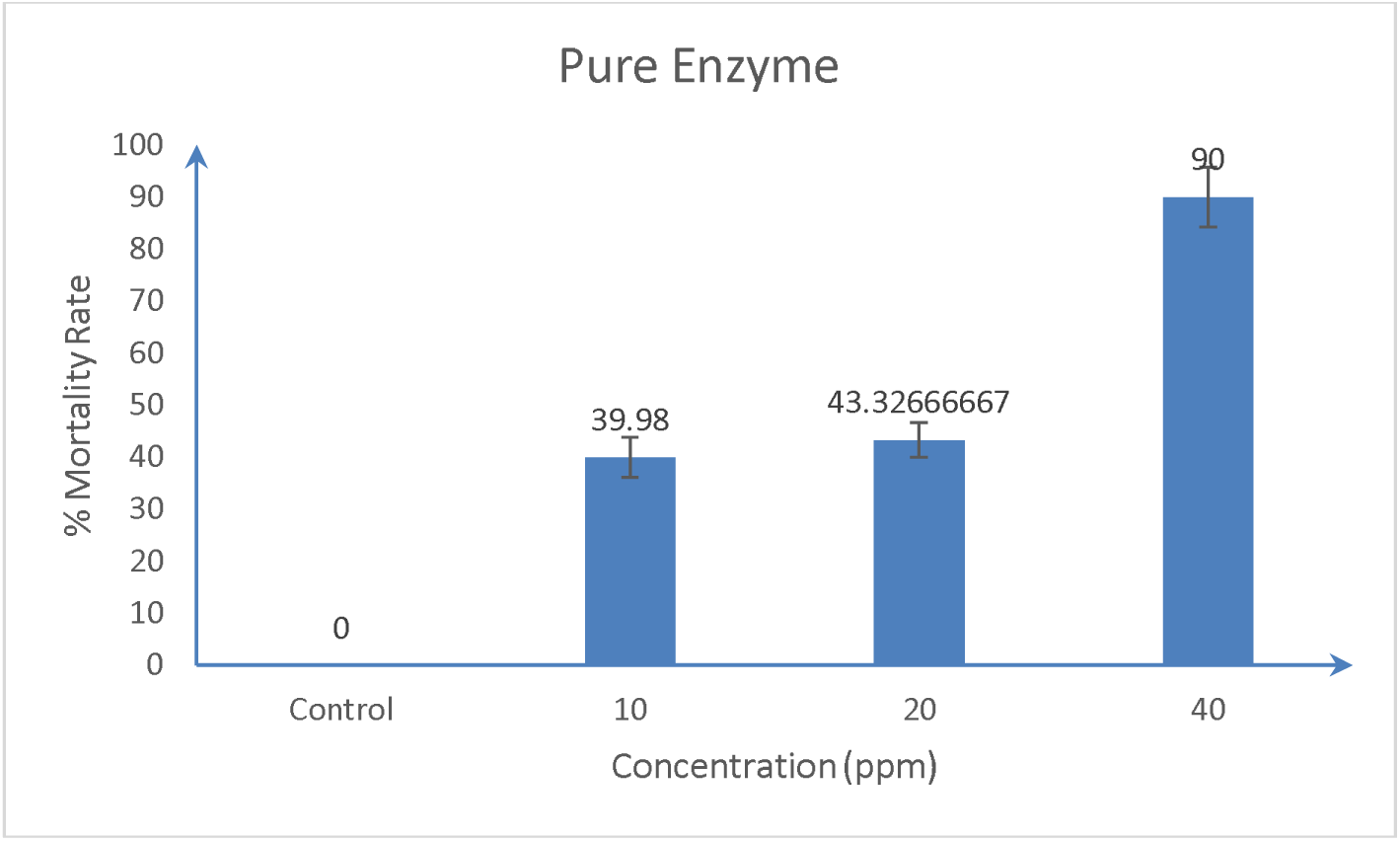
Insecticidal effects of chiA enzyme (crude and pure) on *M. persicae* at different concentrations

#### 3.4.1 ChiA protein similarity and comparison of the results

The amino acid sequence (Supporting Information Fig. S11) and the nucleotide sequence of the chiA protein belonging to the *S. marcescens* GBS19 isolate are depicted in Fig. S12. Chitinase proteins with the closest similarity to the chiA protein were downloaded from the NCBI database, and the phylogenetic relationships between these proteins were analyzed using the Neighbour-Joining method, resulting in the construction of a phylogenetic tree. The results of the study indicate that the chiA protein of *S. marcescens* GBS19 was in specific group (indicated with yellow) within the branches in the phylogenetic tree (Fig. 5.). The Neighbour-Joining analysis revealed that the cloned sequence exhibited a genetic difference of 83/1000 compared to the group branches closest to it. Amino acid data was used to facilitate comparisons with other chitinase proteins in the NCBI database. The results demonstrated that the chiA protein of the isolate exhibited a complete divergence from the other proteins at the 12th, 313th, and 490th amino acid positions (Fig. 6.).

**Figure 5.**
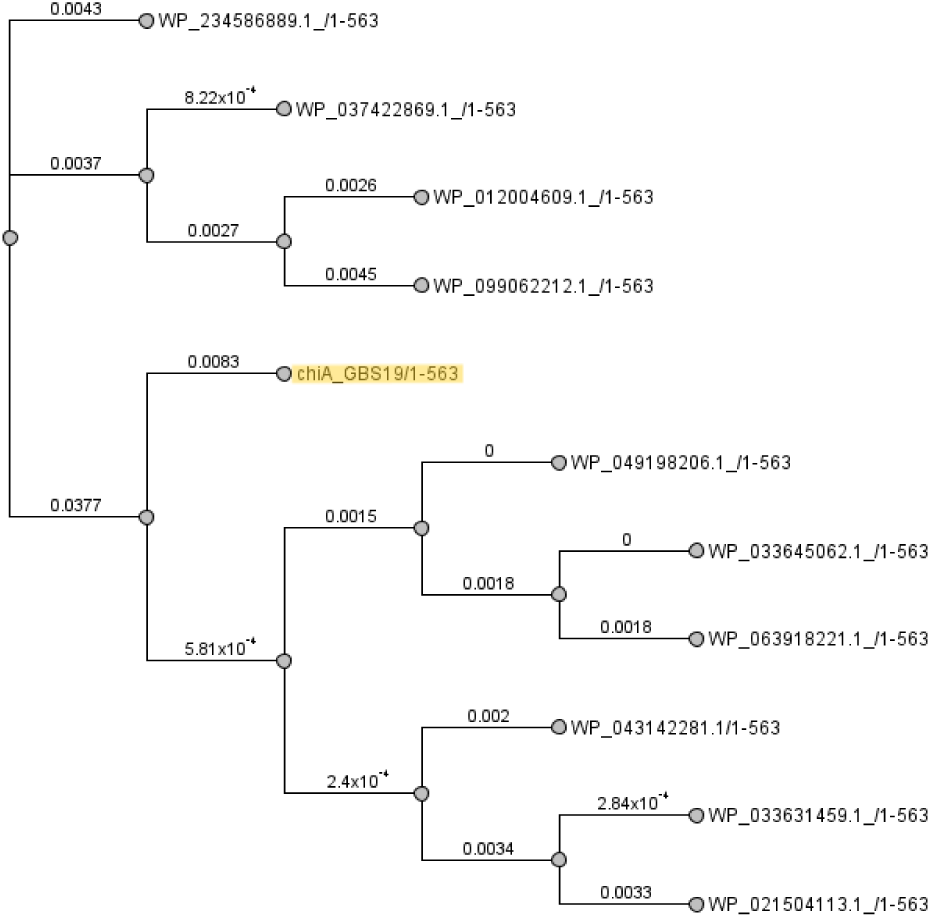
Phylogenetic tree of chiA proteins using Neighbour-Joining

**Figure 6.**
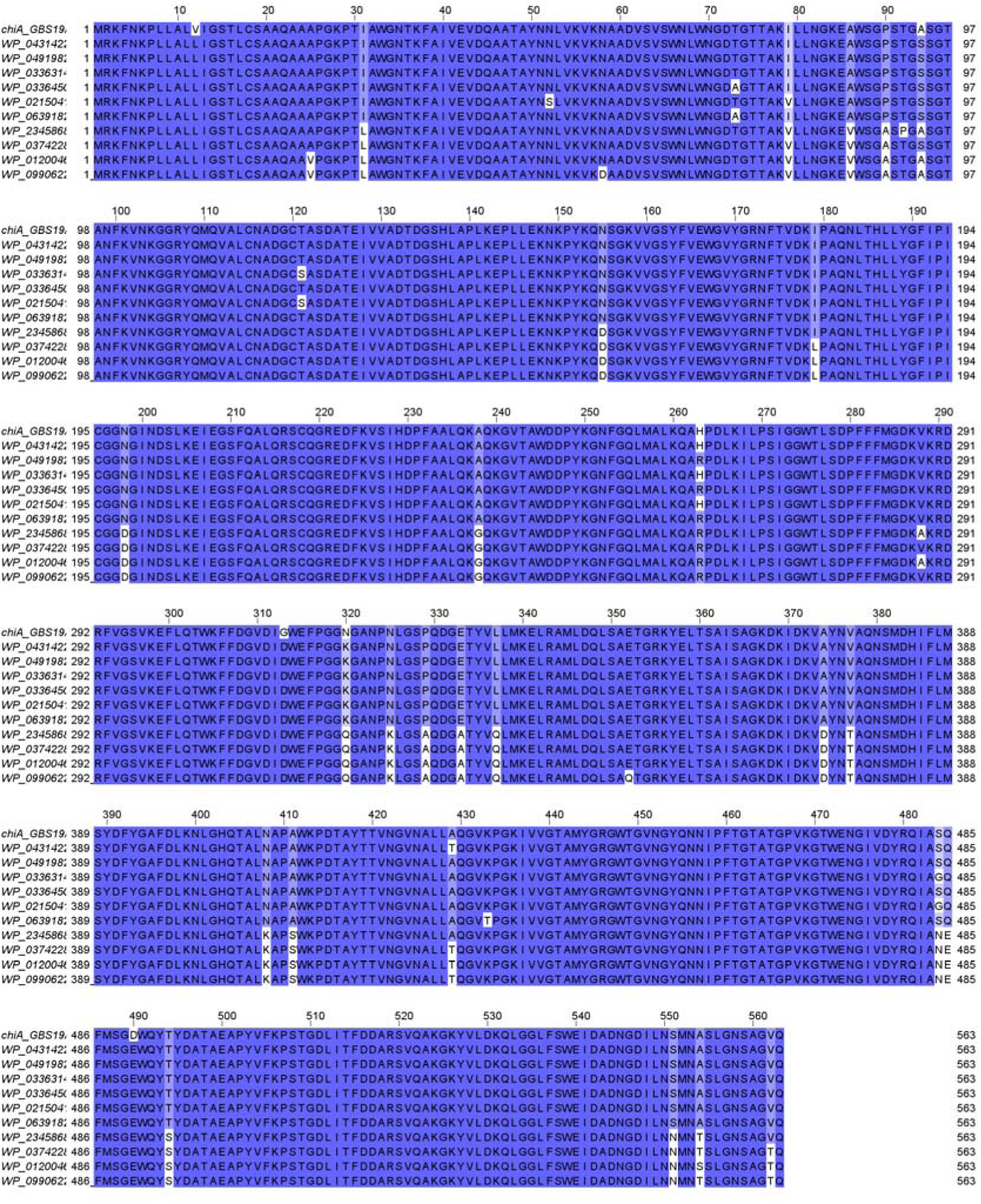
Comparison of ChiA proteins amino acid sequences

#### 3.4.2 ChiA-chitin binding affinity protein- ligand interaction

The results of the analyses indicated that the highest binding affinity between chiA and chitin was -4.10 kcal/mol. In addition, binding occurred at seven different locations on the enzyme, with binding energies varying between -4.10 and -3.63 kcal/mol. The findings regarding the placement results and attachment patterns are given in Fig. 7. The most favourable bonding and the highest affinity interaction points were observed at the following coordinates: TYR390, MET388, ASN391, SER364, ALA365, PHE316 and LYS369 (Supplementary File. S13).Furthermore we visualized the interaction between chitinase enzyme and Chitin layer using modelling program CHARM-GUI according to modified parameters.

**Figure 7.**
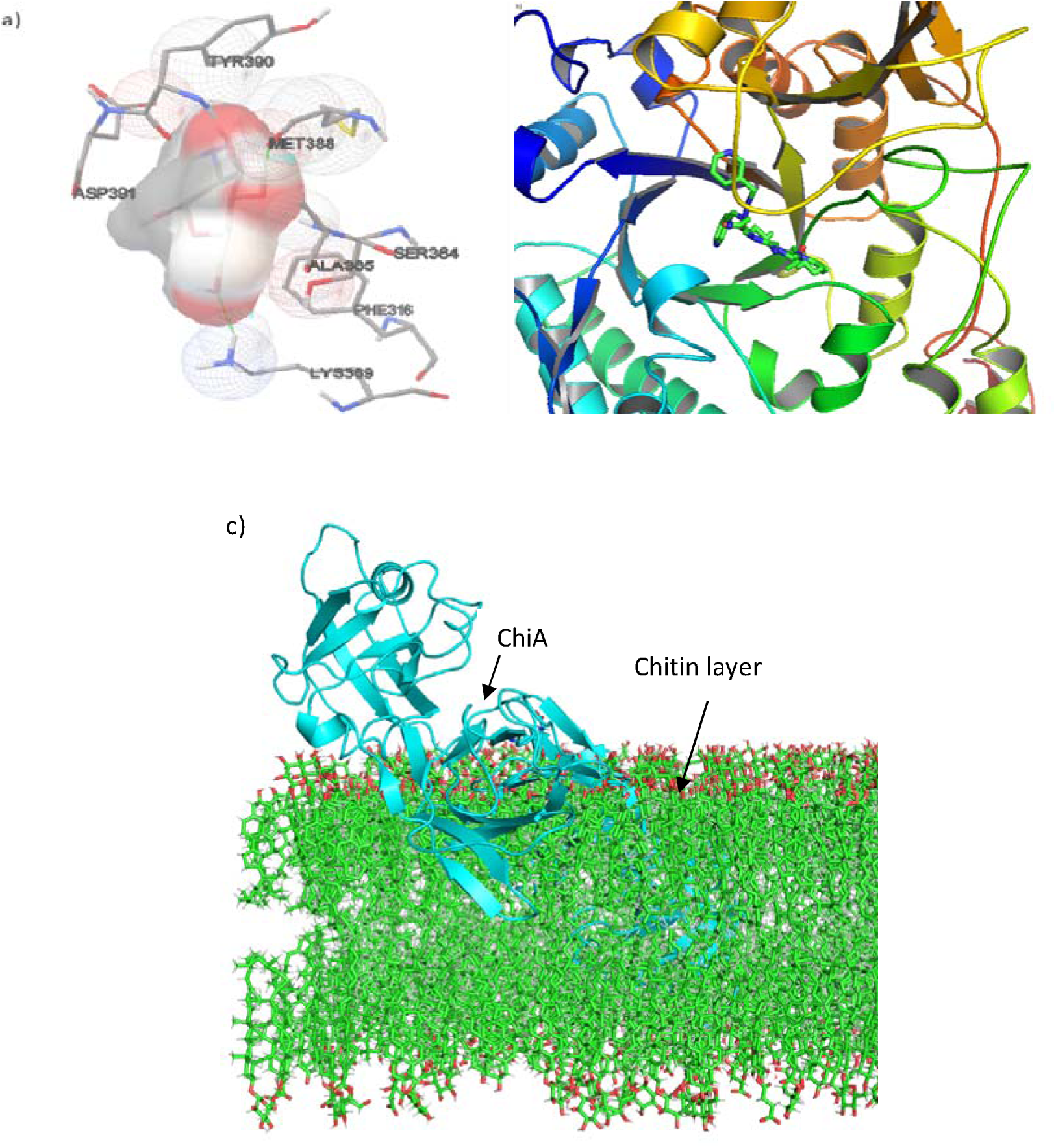
Interaction between chiA protein (PDB 7C92) and chitin substrate and its binding position a) illustration shows the highest binding coordinates b) interaction at highest bonded site c) interaction between chitin layer and chiA protein.

## 4 DISCUSSION

The deleterious effects of the extensive utilization of synthetic pesticides on the environment and human health have prompted the pursuit of environmentally benign techniques for the management of plant diseases and pests. In this context, biocontrol based on the use of beneficial organisms or their products (bioactive molecules or hydrolytic enzymes) holds the greatest promise and are well known as part of integrated pest management. Chitinases are particularly attractive for this purpose because they have fungicidal, insecticidal, and nematicidal activities.^26^ One of the key targets for chitinase enzymes is the chitin polymers found in the intestinal and exoskeleton structures of insects. Consequently, chitinase enzymes produced by microorganisms represent microbial agents with the potential to be employed in the fight against harmful insects.^27^ To date, various chitinase genes were cloned from different bacterial strains such as *Bacillus licheniformis* LHH100, *Chitinibacter* sp. GC72, *Bacillus holodurans* C-125, *Pseudoalteromonas tunicata* CCUG 44952T, *Serratia proteamaculans* 568, *Chromobacterium violaceum* ATCC 12472, *Bacillus altitudinis* KA15, *Paenibacillus barengoltzii*, *Cohnella* sp. A01, *Serratia marcescens* B4A, *Hydrogenophilus hirschii* KB-DZ44, *Xenorhabdus nematophila* HB310 and heterologously expressed.^28,29,30,31,32,33,34,35,36,37,38,39^

Among gram (-) negative bacteria, *S. marcescens* is considered one of the most efficient degraders of chitin in nature and is used as a model system to study chitinase functionality.^40^ In this study, the GBS19 (Assembly Number: GCA_021474915) strain belonging to the *S. marcescens* species, isolated from soil and with a complete genome sequence, was used. The chiA gene from *S. marcescens* is known to degrade β-chitin much more efficiently than chiB and chiC. A comparable observation was made regarding the degradation of α-chitin. The results of these data indicate that the high activity of chiA is due to its capacity to access crystalline substrate to a greater extent than its catalytic efficiency.^41^

To replicate the chiA gene region present in the *S. marcescens* GBS19, gene-specific primers have been designed and amplified via polymerase chain reaction (PCR). The amplified gene region was subsequently cloned into the pBluescript II KS(+) vector. To find the accuracy of the cloned gene region, the clones were subjected to next-generation sequencing (NGS) sequence analysis and the results were evaluated. Upon comparison of the sequence results received from the company with the sequence of the existing gene region, it was confirmed that the 1692 bp chiA gene region was successfully cloned into the vector without base loss or change. Therefore, the chiA protein of our isolate was compared with the reference sequences of chitinase proteins in the NCBI database. The results of the comparison indicated that the chiA protein of our isolate exhibited 98.93% similarity to the *S. marcescens* chitinase protein (WP_043142281) in the database, 98.76% similarity to the *S. nematodiphila* chitinase protein (WP_033631459), 98.58% similarity to the *S. surfactantfaciens* chitinase protein (WP_063918221), and 98.58% similarity to the *S. entomophila* chitinase protein (WP_063918221) in the database. The protein in question (WP_234586889) exhibited a 94.85% similarity to the chitinase protein of *S. grimesii* (WP_037422869).

Horn et al. (2006) successfully cloned and expressed the chitinases of the *S. marcescens* BJL200 strain using *E. coli* as the host organism.^42^ Similarly, Suzuki et al. (2002) and Hult et al. (2005) selected the *E. coli* host organism for the cloning and expression of chitinases from the *S. marcescens* 2170 isolate in their study.^41,43^ The cloning phase of this study was conducted using *E. coli* TOP10 as the host organism, while the expression phase employed *E. coli* BL21(DE3). This approach was informed by similar studies reviewed in the literature, which had achieved fruitful results. Following the cloning phase, the chitinase gene region was cloned into the pET-22b (+) vector with the T7 RNA polymerase promoter, and protein expression was performed in *E. coli* BL21(DE3) bacteria. The expressed chiA protein was purified using Ni-NTA affinity chromatography, with the aid of the his-taq tag present within the vector. The size of the purified chiA protein, including his-taq tail, was confirmed by SDS-PAGE analysis to be 62.5 kDa.

Following the measurement of the activity of the pure chiA enzyme, the optimum pH and temperature values at which the enzyme operated were calculated. No significant differences were observed in the measurements made in the pH range of 3-10. However, it was noted that the enzyme activity was measured at a higher level at pH 5. In a similar study, Okay and Alshehri (2020) found that the optimum pH for the chiA protein they isolated from *S. marcescens* Bn10 was 9 (alkaline).^44^ Likewise, Aggarwal et al. (2016) reported the optimum pH of the chitinase protein they isolated from *S. marcescens* as 9.^45^ Similarly, Danışmanzoğlu et al. (2015) isolated chiA from *S. marcescens* and found the optimum pH to be 8.^46^ Consequently, Babashpour et al. (2012) determined that the optimum pH of chiA, which they purified from *S. marcescens* B4A, was 6 (acidic).^37^ Nawani and Kapadnis (2001) demonstrated that the chitinase from the *S. marcescens* NK1 isolate exhibited the highest activity at pH 6.2.^47^ In a separate study, Li et al. (2020) identified the optimal pH for the chiA enzyme purified from *S. marcescens* as 6.^48^ Zarei et al. (2011) employed the *S. marcescens* B4A isolate in their study, yet the optimal pH of the chiA enzyme was established by Babashpour et al. (2012), who determined the pH to be 5, which differs from their findings.^37,49^ Moon et al. (2017) determined that the optimum pH of the chitinase enzyme isolated from *S. marcescens* PRNK-1 was 5.5.^50^ A review of the literature revealed that the optimal pH for the chiA enzyme in different isolates of *S. marcescens* bacteria may vary.

The optimal temperature for the chiA enzyme of the soil-origin *S. marcescens* GBS19 isolate was determined to be 40°C. “Li et al. (2020) determined the optimum temperature of the chiA enzyme and purified chiA from Serratia marcescens obtained from Lentinula edodes (shiitake mushroom), commonly used in Far Eastern cuisine . Additionally, Okay and Alshehri (2020) investigated the S. marcescens chiA protein, which was isolated from Balaninus nucum (hazelnut worm), and reported its optimum temperature as 60°C . The optimal temperature for the chiA enzyme in this bacterium was 37°C . A review of the literature indicates that chitinases isolated from *S. marcescens* bacteria of different origins exhibited optimal activity at varying temperatures. The optimal temperature of the chiA enzyme, as identified in this study, closely resembles the ambient air temperatures commonly observed in regions where the agricultural pest peach aphid thrives during the spring and summer seasons. This suggests a potential application of the enzyme in biological control. In their study, Broadway et al. (1998) demonstrated that endo-chitinases belonging to the *Streptomyces albidoflavus* bacterium significantly reduced the survival rate of the insect when applied against the pest *M. persicae*.^51^ Danışmanzoğlu et al. (2015) reported an insecticidal activity of chiA isolated from *S. marcescens* against *H. armigera* larvae.^46^ Furthermore, it has been demonstrated that a chitinase purified from *B. laterosporus* exhibits insecticidal activity against *Plutella xylostella* (cabbage leaf moth).^52^

Various sources have been explored for chitinase production, leading to the discovery of numerous chitinases with enhanced properties. However, naturally produced chitinases are not suitable for industrial applications due to their incompatibility with industrial formats. Genetic engineering has therefore served as an intermediary between industrial and natural strains. A notable increase in the efficiency of chitinases has been observed in recombinant organisms compared to wild strain.^53^ The applications of chitinases are well documented, with a particular focus on their role as biocontrol agents in various fields, including agriculture. The development of recombinant enzyme production systems has emerged as a promising platform for the efficient industrial production of chitinase.^8^ Microbial chitinases have demonstrated their potential to produce large quantities of various chitinases.^54^ Given the multitude of applications for chitinases; it is anticipated that demand will increase soon.^55^

It is possible that the use of chitinase enzyme in the study may result in an increase in product yield and a reduction in product losses. Chitinases have the potential to be employed as a biologically based control method against plant pests, as they can break down the insect’s chitin layer, thereby reducing the insect’s nutrition and defence. Concurrently, the utilisation of chitinase enzyme in the pharmaceuticals employed to combat plant pests offers the potential for enhanced production and efficacy per unit area. The global and domestic production of chitinase enzyme is currently limited. The results obtained constituted the foundation for future studies investigating the effects of microbial-derived *S. marcescens* GBS19 chiA protein on *M. persicae*. Indeed, further research is essential to explore the insecticidal properties of the chiA protein and identify the specific insect species suitable for effective pest control. The study established the potential of the enzyme in question for use as a biopesticide. Further investigations will delineate the potential of this enzyme for industrial and field applications.

Although any biological agent has genetic properties that are harmful to humans and the environment, we have the chance to go beyond these limits with recombinant DNA technology, which we emphasize in our article. We anticipate that this study will pave the way as a pioneering investigation in this field for the future.

## ACKNOWLEDGEMENTS

This study was supported by Muğla Sıtkı Koçman University Scientific Research Projects Coordination Unit with project number 20/114/02/1/5. We would like to thank the Scientific Research Coordination Unit for their support. At the same time, this study has been supported by TUBİTAK 2210-C Priority Areas with the application number ’’ 1649B02203359 ’’ of the Domestic Master Scholarship Program. We would like to thank TUBİTAK for their support. The authors declare that whole studies have been carried out in the frame of Ahmet Can’s M.Sc. thesis under the supervision of Ö. Baysal (Ph.D. Professor in Molecular Microbiology)

## AUTHOR CONTRIBUTIONS

Ahmet Can: Conceptualization, Data curation, Formal analysis, Investigation, Methodology, Project administration, Resources, Software, Validation, Visualization, Writing – original draft, Writing – review & editing. Ömür Baysal: Conceptualization, Data curation, Formal analysis, Funding acquisition, Investigation, Methodology, Resources, Software, Supervision, Validation, Visualization, Writing – original draft.

